# Characterization of a novel NDM-5-harboring plasmid from a carbapenem-resistant *Escherichia coli* isolate from China

**DOI:** 10.1101/2020.01.24.919274

**Authors:** Dongdong Yin, Yanfeng Lin, Zhonghong Li, Hui Ma, Lanfen Lu, Kaiying Wang, Lang Yang, Xinying Du, Hongbin Song, Peng Li, Kezong Qi

## Abstract

A carbapenem-resistant *Escherichia coli* (sequence type 5415) strain was isolated from a male patient. S1-PFGE and Southern blotting showed that the *bla*_NDM-5_ gene was located on a novel 66-kb IncFII [F2:A-:B-] plasmid. Conjugation assays revealed that the *bla*_NDM-5_-bearing plasmid was self-transferrable. Genomic sequencing and comparative analysis suggested that plasmid p2947-NDM5 likely originated from a combination of an IncFII-type backbone and the *bla*_NDM-5_ flanking genetic elements. This study highlights the genetic complexity of an NDM-5 carbapenemase and the urgent need for continuous active monitoring.

**IMPORTANCE:** Carbapenem resistance due to the NDM enzyme has become a serious clinical problem in medical care and public health. We here identified a *bla*_NDM-5_-positive *E. coli* strain ST5414. The *bla*_NDM-5_ gene was located on a novel self-transferrable IncFII type plasmid p2947-NDM5. Genomic sequencing and comparative analysis suggested that plasmid p2947-NDM5 likely originated from a combination of an IncFII-type backbone and the blaNDM-5 flanking genetic elements. The genetic context of NDM-5 is highly similar to that of animal origins, and we speculate that the *bla*NDM-harboring IncFII plasmid has a potential risk of being transmitted from animals to humans and should be paid more attention. Our findings highlight the threat of NDM-5 carbapenemase circulation and the urgency of stringent surveillance and control measures.

---

Carbapenemase-producing *Enterobacteriaceae* (CPE) constitute a public health problem in terms of both hospital- and community-acquired infections(1). Since the identification of NDM-1 in a Swedish traveler returning from India(2), NDM enzymes have received special attention due to their rapid global spread and frequent association with other resistance genes. NDM-5 was first identified in an *Escherichia coli* strain isolated from a patient who had been hospitalized in India(3). Since then, NDM-5 has been detected in different countries around the world. The amino acid sequence of NDM-5 differs from that of NDM-1 at positions 88 (Val→Leu) and 154 (Met→Leu), which confer a high level of hydrolytic activity against carbapenems(3). The rapid evolution and dissemination of NDM-5 represent a crucial challenge for clinical treatments.

Extraintestinal *E. coli* is a relatively common pathogen causing community and health care-related infections among *Enterobacteriaceae* in China. The acquisition of NDM is a great concern since it would greatly limit the treatments for *E. coli* that frequently carry multiple resistance determinants. Identifying clones or plasmids with *bla*_NDM_ genes is important for understanding the epidemiology of resistance and controlling the spread of NDM in communities and healthcare systems(4).

In this study, we report the emergence of the NDM-5-producing *E. coli* strain ST5414 in China and characterized a novel plasmid carrying the *bla*_NDM-5_ gene using Illumina and Nanopore sequencing platforms.

## RESULTS

### Bacterial identification and Susceptibility Testing

Strain ECO2947 was identified as *E. coli* using the Vitek 2 compact system and confirmed by 16S rRNA sequencing. The MIC values of the tested antimicrobials revealed that *E. coli* ECO2947 exhibited resistance to nearly all tested β-lactam antibiotics, including ampicillin, sulbactam/ampicillin, piperacillin/tazobactam, ceftriaxone, ceftazidime, cefepime, cefamedin and imipenem, with the exception of aztreonam (Table 1). PCR amplification and sequencing confirmed the presence of *bla*_NDM-5_.

**TABLE 1.** Antibiotic susceptibilities of *E. coli* ECO2947 and the *E. coli* J53 transconjugants.

| Antimicrobial | MIC (µg/ml) |  |  |
| --- | --- | --- | --- |
|  | ECO2947 | J53 (the transconjugant) | J53 |
| Ampicillin | ≥32 | ≥32 | 8 |
| Sulbactam-Ampicillin | ≥32 | ≥32 | 4 |
| Aztreonam | ≤1 | ≤1 | ≤1 |
| Furadantin | 32 | ≤16 | ≤16 |
| Ciprofloxacin | ≤0.25 | ≤0.25 | ≤0.25 |
| Piperacillin-Tazobactam | 64 | 64 | ≤4 |
| Gentamicin | ≤1 | ≤1 | ≤1 |
| Cefepime | 16 | 8 | ≤1 |
| Ceftriaxone | ≥64 | ≥64 | ≤1 |
| Ceftazidime | ≥64 | ≥64 | ≤1 |
| Cefotetan | 32 | 32 | ≤4 |
| Cefamedin | ≥64 | ≥64 | ≤4 |
| Tobramycin | ≤1 | ≤1 | ≤1 |
| Imipenem | ≥16 | ≥16 | ≤1 |
| Amikacin | ≤2 | ≤2 | ≤2 |
| Levofloxacin | 1 | ≤0.25 | ≤0.25 |

### Microbiological and Genomic Features of *E. coli* ECO2947

S1 PFGE showed that *E. coli* ECO2947 contained two different plasmids (~66 kb and ~108 kb) (Figure 1). Southern blotting revealed that the *bla*_NDM-5_ gene was located on the ~66 kb plasmid (named p2947-NDM5), which was transferred to *E. coli* J53 at a frequency of 1.63×10^-2^ transconjugants per donor cell.

**FIGURE 1.**
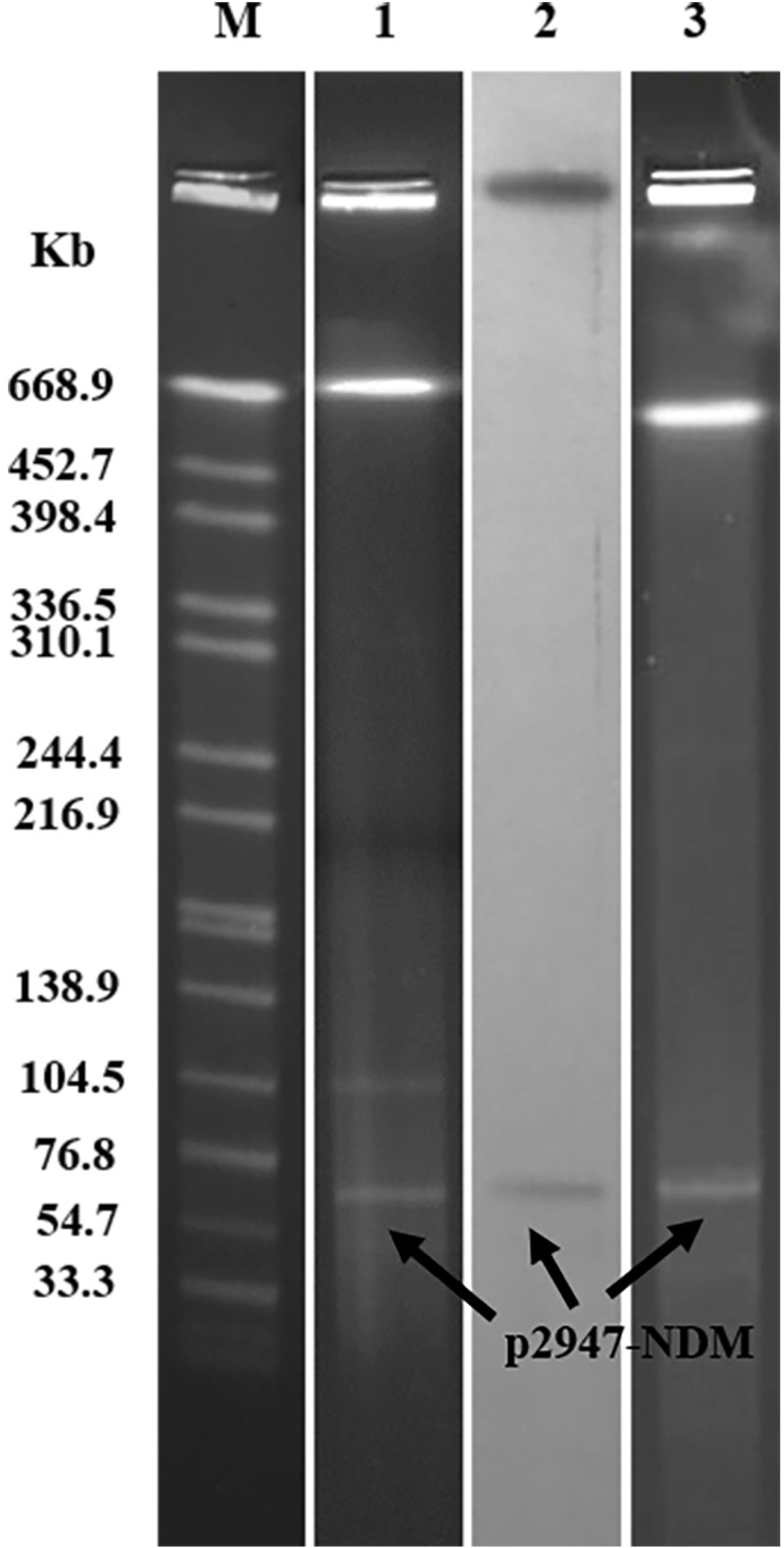
S1-PFGE pattern for strain ECO2947 and Southern blotting for the *bla*_NDM-5_ gene. Lanes: Marker, *Salmonella* serotype Braenderup strain H9812 as the size standard; 1, PFGE result for S1-digested plasmid DNA of strain ECO2947; 2, Southern blotting with the probe specific to *bla*_NDM-5_; 3, PFGE patterns for S1-digested plasmid DNA of *E. coli* transconjugants J53.

The transconjugants acquired resistance to ampicillin, ceftriaxone, sulbactam/ampicillin, piperacillin/tazobactam, ceftazidime, cefamedin, and imipenem (Table 1), and the MIC values of carbapenems against the transconjugants were considerably increased compared with that against the recipient strain *E. coli* J53.

*E. coli* ECO2947 was further subjected to whole genome sequencing using both MiSeq and MinION sequencing. Genomic analysis revealed that strain *E. coli* ECO2947 belonged to a novel sequence type ST5414 and had a 4,884,967 bp chromosome and two plasmids. Twelve virulence factors were found in the genome: single copies of eae (intimin), espA (type III secretion system), espB (secreted protein B), espF (type III secretion system), iss (increased serum survival), lpfA (long polar fimbriae), nleA (non-LEE encoded effector A), nleB (non-LEE encoded effector B), nleC (non-LEE encoded effector C), tir (translocated intimin receptor protein) and two copies of gad (glutamate decarboxylase). A screening for acquired resistance determinants found that the chromosome only possessed the resistance gene *mdf(A)*, while the plasmid p2947-NDM5 carried only *bla*_NDM-5_, and the other plasmid (named p2947-D) carried multiple resistance genes, including *sul2, qnrS1, aph(3”)-Ib* and *aph(6)-Id*.

### Characterization of the novel blaNDM-5-harboring plasmid

The *bla*_NDM-5_-harboring plasmid p2947-NDM5 belonged to the incompatibility type IncFII [F2:A-:B-] with a length of 66,053 bp, an average G + C content of 52.41% and 94 predicted coding sequences. p2947-NDM5 had a 61-kb backbone and a 5-kb multidrug resistance (MDR) region. A BLAST search revealed that p2947-NDM5 was highly similar to plasmid p974-NDM of *E. coli* strain 974 (99% coverage and 99.66% identity), plasmid unnamed4 of *Klebsiella pneumoniae* strain 4743 (93% coverage and 100% identity, referred to as p4743) and the plasmid of *Salmonella enterica* subsp. *enterica* serovar Derby strain 75 (92% coverage and 99.97% identity, referred to as p75). The four plasmids have almost identical backbones and contain a set of core genes responsible for plasmid replication (*repA*), conjugation/T4SS (*tra* and *trb* genes), stability (*stdB*) and segregation (*parM*) (Figure 2A). However, p2947-NDM5 had quite different MDR regions from these similar plasmids. Plasmid p2947-NDM5 had an MDR region organized as IS*5-bla*_NDM-5_-*ble*_MBL_-*trpF-tat*-Δ*ctuA1*-IS*26*, while p974-NDM had the *bla*_NDM-1_ gene instead and an additional region composed of IS*26*, ISKox*3*, resolvase, Tn2 transposon, IS*3000* and ΔIS*Aba125*. Plasmids p75 and p4743 had quite different resistance genes and genetic contexts. Both p75 and p4743 had two copies of IS*26* at each end; the former was composed of IS*26*-IS*903*-BtuB-tnpR Tn*3*-IS*26*, while the latter was composed of IS*26*-intl*1*-IS*26*-Δ*bla*_TEM_-*WbuC-bla*_CTX-M-15_-IS*26* (Figure 2B).

**FIGURE 2.**
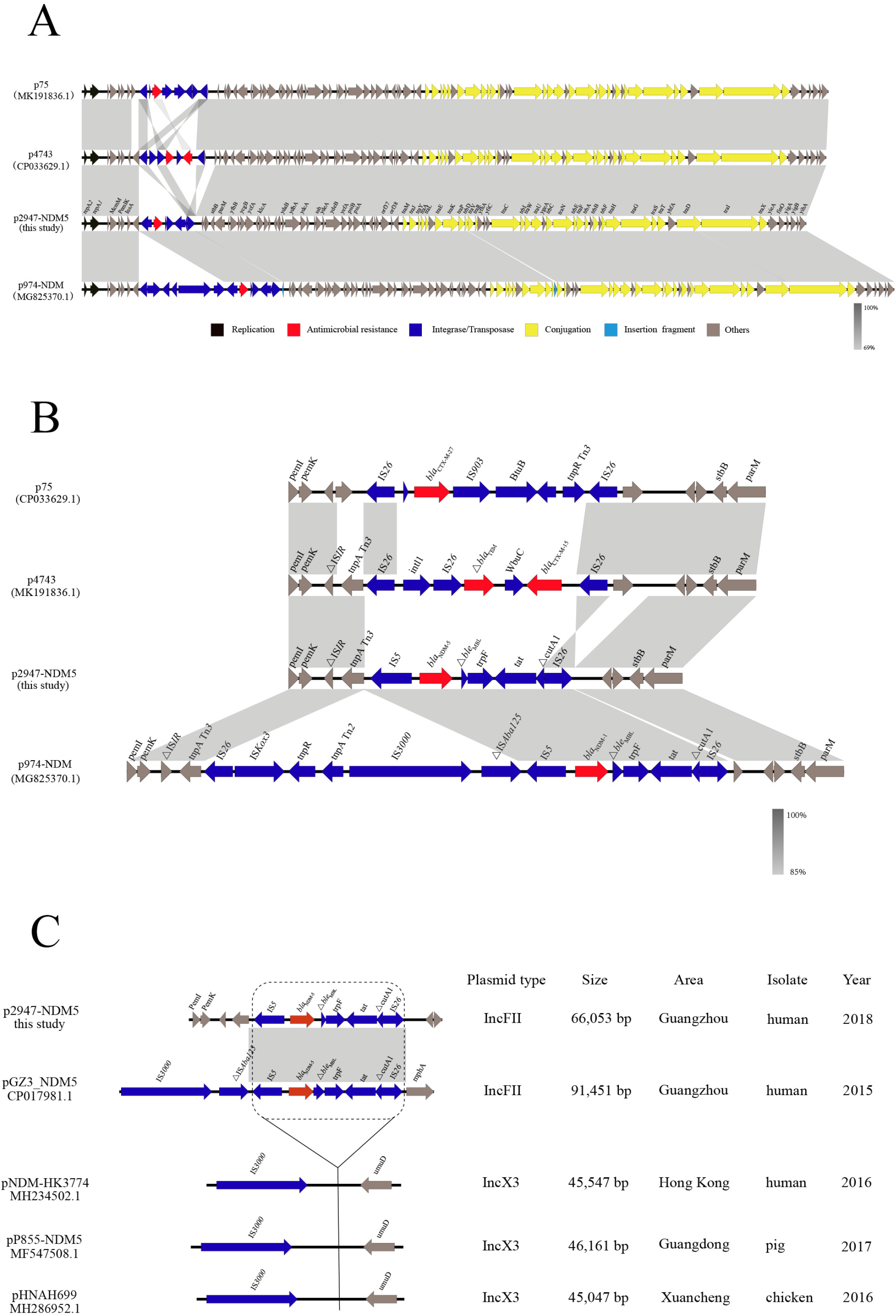
Plasmid analysis of p2947-NDM5. (A) Genetic structure comparison of p2947-NDM5, p75(CP033629.1), p4743(MK191836.1) and p974-NDM(MG825370.1). (B) Comparative analysis of MDR regions.(C) Comparative analysis of the genetic contexts of *bla*_NDM-5_ in plasmids reported in this study and previously described.

Comparative analysis revealed that the genetic context of *bla*_NDM-5_ (~5,100 bp) in p2947-NDM5 was nearly identical to those previously reported in pGZ3-NDM5 (100% coverage and 100% identity), pNDM-HK3774 (100% coverage and 100% identity), pHNAH699 (100% coverage and 100% identity) and pP855-NDM5 (100% coverage and 99.98% identity) (Figure 3C).

## DISCUSSION

Due to the intrinsic and acquired resistance of *E. coli*, this bacterium constitutes a serious clinical threat that limits the choice of treatment. Most reports have indicated a high ST diversity for *bla*NDM-5-positive *E. coli*(5-7). The MLST analysis revealed that *E. coli* ECO2947 belongs to ST5414, which is unlike the ST types of NDM-5-producing *E. coli* ST167 (China)(6), ST540 (Japan)(8), ST648 (Australia), ST648 (UK)(3) and ST648 (India). This appears to be the first report of an ST5414 *E. coli* strain expressing an NDM-5 β-lactamase. This is a worrying development, as it demonstrates the further spread of *bla*_NDM-5_ among different ST types of *E. coli,* and the transfer frequency of p2947-NDM5 demonstrated its great potential to transfer across species.

Horizontal gene transfer also promotes the widespread dissemination of *bla*_NDM_ in *Enterobacterales*(9). Animals have been considered a potential reservoir of MDR gram-negative organisms due to the extensive use of antibiotics in animals as growth promoters and for treatment purposes(10). Investigations in recent years confirmed the endemic presence of ESBL-producing *E. coli* isolates in major food-producing animals (chickens, pigs and cattle) with increasing prevalence(11, 12), and integron plasmid vehicles in animal isolates were identical or highly similar to those carried by clinical *E. coli* strains isolated from asymptomatic humans(13). NDM gene transfer carried by food animals will lead to the spread of the gene along the food chain, ultimately causing harm to human health and public health(14, 15). *Klebsiella pneumoniae* strain 4743 was isolated from humans in Italy, the *Salmonella enterica* subsp. *enterica* serovar Derby strain was isolated from swine in the United States, and the *E. coli* strain 974 was isolated from a pig in Hong Kong. In this study, two of the three plasmids highly similar to p2947-NDM5 were from animals. Additionally, the genetic context of *bla*_NDM-5_ of p2947-NDM5 has also been found on different types of plasmids and in different species. Although the occupation of the patient is unknown, it is still possible that the patient was in contact with animals, and a conjugation assay revealed that p2947-NDM5 was self-transferrable. From this, we speculated that p2947-NDM5 was the genetic context of *bla*_NDM-5_, which had been inserted into the IncFII [F2:A-:B-] plasmid backbone during transmission, and the *bla*_NDM_-harboring IncFII plasmid has a potential risk of being transmitted from animals to humans through the food chain. Therefore, the association of IncFII plasmids and *bla*_NDM_ variants and the epidemiology of IncFII plasmids in Enterobacteriaceae warrant more studies.

## CONCLUSION

In summary, we identified a *bla*_NDM-5_-positive *E. coli* strain, ST5414, for the first time. The *bla*_NDM-5_ gene was located on a novel self-transferrable IncFII-type plasmid. Our study highlights the potential spread of carbapenem-resistant plasmids among *Enterobacteriaceae*. Further research is necessary to take urgent and effective surveillance measures and to control the spread of the *bla*_NDM-5_-carrying IncFII plasmids.

## MATERIALS AND METHODS

### Identification of the *E. coli* Strain Carrying bla_NDM_

A carbapenem-resistant strain, ECO2947, was recovered from a sputum culture of a patient through routine surveillance in 2018 in Guangzhou, China. The species of strain ECO2947 was identified by the Vitek 2 compact system (bioMérieux, France) and confirmed by amplification and sequencing of the 16S rRNA gene(16). The presence of genes encoding carbapenemases and ESBLs was determined by PCR and sequencing(17).

### S1-PFGE, Southern blotting and Conjugation

Bacterial genomic DNA from strain ECO2947 was prepared in agarose plugs and digested with the S1 endonuclease (Takara, Dalian, China). DNA fragments were separated by pulsed-field gel electrophoresis (PFGE) through a CHEF-DR III system (Bio-Rad, Hercules, USA). The plasmid DNA was transferred to a positively charged nylon membrane (Solabio, China) and hybridized with the digoxigenin-labeled specific probe to *bla*_NDM-5_. Conjugation experiments were performed by broth and filter mating using strain ECO2947 as the donor and azide-resistant *E. coli* J53 as the recipient. Strains ECO2947 and J53 were mixed (ratio of 1:3) in Luria-Bertani (LB) broth, which was used to make LB agar plates, and incubated for 18 h at 37 °C. The mixture was spread on a selective MacConkey agar plate containing meropenem (4 μg/ml) and sodium azide (150 μg/ml) to select transconjugants. Horizontal transferability of drug resistance was evaluated by antimicrobial susceptibility testing, and the corresponding transconjugants were confirmed by S1-PFGE.

### Antimicrobial Susceptibility Testing

The minimal inhibitory concentrations (MICs) of amikacin, ampicillin, sulbactam/ampicillin, aztreonam, furadantin, ciprofloxacin, piperacillin/tazobactam, gentamicin, cefepime, ceftriaxone, ceftazidime, cefotetan, cefamedin, tobramycin, imipenem and levofloxacin were determined by a Vitek 2 compact system (BioMérieux, France) following the manufacturer’s instructions. The results were interpreted following the guidelines of the Clinical and Laboratory Standards Institute (CLSI)(18).

### Whole Genome Sequencing and Analysis

Genomic DNA was extracted using a High Pure PCR Template Preparation Kit (Roche, Basel, Switzerland). Whole genome sequencing was carried out using both Illumina MiSeq and Oxford Nanopore MinION. The *de novo* hybrid assembly of short Illumina reads and long MinION reads was performed using Unicycler v0.4.8(19) with the conservative mode. Complete circular contigs were corrected using Pilon with Illumina reads for several rounds until no change was detected. Genome sequences were annotated using the RAST server(20). The sequence type was determined through the MLST web server(21). Virulence genes and plasmid types were identified using VirulenceFinder, PlasmidFinder and pMLST(22).

### Nucleotide Sequence Accession Number

The complete sequences of the chromosome of strain ECO2947, plasmid p2947-D and p2947-NDM5 have been deposited in GenBank under accession numbers CP046259, CP046260 and CP046261, respectively.

## ACKNOWLEDGMENTS

The study was supported by grants from the National Science and Technology Major Project (no. 2018ZX10201001-003 and no. 2018ZX10712001-002-002), the Beijing Natural Science Foundation (no. 5172029) and the Beijing Noval Program (no. Z181100006218110).

## Footnotes

1 *Dongdong Yin, Yanfeng Lin, Zhonghong Li and Hui Ma contributed equally to this work. Author order was determined by the relative amount of contributions*.

